# A simple strategy for sample annotation error detection in cytometry datasets

**DOI:** 10.1101/2021.10.26.465993

**Authors:** Megan E Smithmyer, Alice E Wiedeman, David A.G. Skibinski, Adam K. Savage, Carolina Acosta-Vega, Sheila Scheiding, Vivian H. Gersuk, S. Alice Long, Jane H. Buckner, Cate Speake

## Abstract

Mislabeling samples or data with the wrong participant information can impact study integrity and lead investigators to draw inaccurate conclusions. Quality control to prevent these types of errors is commonly embedded into the analysis of genomic datasets, but a similar identification strategy is not standard for cytometric data. Here, we present a method for detecting sample identification errors in cytometric data using expression of HLA class I alleles. We measured HLA-A*02 and HLA-B*07 expression in 3 longitudinal samples from 41 participants using a 33-marker CyTOF panel designed to identify major immune cell types. 3/123 samples (2.4%) showed HLA allele expression that did not match their longitudinal pairs. Furthermore, these same three samples’ cytometric signature did not match qPCR HLA class I allele data, suggesting that they were accurately identified as mismatches. We conclude that this technique is useful for detecting sample labeling errors in cytometric analyses of longitudinal data. This technique could also be used in conjunction with another method, like GWAS or PCR, to detect errors in cross-sectional data. We suggest widespread adoption of this or similar techniques will improve the quality of clinical studies that utilize cytometry.

## INTRODUCTION

Sample annotation errors occur when clinical trial samples are paired with the incorrect participant metadata. This type of error can produce misleading results which can negatively impact reproducibility, and even undermine our understanding of biological phenomena. Investigators are increasingly aware of sample annotation errors, as evidence suggests they occur in many publicly available datasets (1-5). The reported rates of incorrect sample annotation vary. One study found 40% of publicly available datasets investigated contained at least one such error (2). Others reported that roughly 2% (6) or 3% (1) of all samples analyzed, including samples from a global network of biobanks (6) or publicly available genomic data from multiple studies (1), were paired with the wrong metadata. The high prevalence of these errors, and the recognition that even a small number of errors can negatively impact study integrity, has led to the adoption of quality control protocols designed to detect problematic samples in genomic and transcriptomic studies (1, 6-9). However, the causes of sample misannotation are not limited to genomic analysis. These types of issues can also occur in cytometry datasets, but there are no established methods for the molecular identification of a sample from information captured within cytometry data. In this communication, we present a simple and generalizable quality control method to detect sample mix-ups in cytometry datasets.

In genomic data, sex prediction based on the presence of genes associated with sex chromosomes has been used to verify participant metadata for over a decade (7). While this approach has several limitations including the inability to identify sample mix-ups that occur between participants of the same sex and potential false positives in participants with sex chromosome anomalies, it has been used to successfully identify annotation errors in publicly available datasets (2). Another widely implemented method utilizes single nucleotide polymorphisms (SNPs) to identify samples collected from the same individual, allowing investigators to detect a larger proportion of errors in longitudinal datasets (1, 8, 9).

Unlike genomic data, cytometry data generally presents a snapshot of a dynamic distribution of cell populations as opposed to an inheritable, immutable characteristic like genotype. This presents a challenge in the identification of individual sample donors, because few characteristics of the sample are sufficiently unique to individual participants and remain unchanged in response to immune perturbations. One solution to this problem is the measurement of highly polymorphic cellular proteins such as human leukocyte antigen (HLA) class I or class II proteins. Because HLA vary greatly among individuals, identifying the presence or absence of specific HLA alleles can help distinguish samples from individuals who do not share the same genotype and can help identify longitudinal samples from the same individual. In fact, the measurement of specific HLA alleles has been used successfully in medical diagnostics to confirm that biopsies are matched with the correct patient (10).

For this study, we successfully integrated antibodies specific for HLA-A*02 and HLA-B*07 into a 33-marker mass cytometry panel that was developed for the identification and characterization of major immune cell populations in whole blood. This approach was used to analyze 3 longitudinal samples collected from each of 16 COVID-19 patients and 25 healthy controls for a total of 123 samples. We compared each individual’s longitudinal HLA measurements by CyTOF to identify sample swaps that occurred at any timepoint for an individual. We also utilize genomic HLA data from one timepoint as an external reference to confirm the HLA type for participants with mismatching longitudinal data. Here, we show that this approach can identify samples with potential annotation errors that can occur during a high throughput immunomonitoring study, facilitating their exclusion from analysis and interpretation of the data.

## MATERIALS AND METHODS

### Study Population

The participants included in this analysis were enrolled in two observational cohorts; 16 (39.0%) were enrolled in a study of COVID-19 positive individuals (11). 25 subjects (61.0%) were enrolled in an investigation of the immune systems of individuals with no personal history of chronic disease, autoimmunity, or severe allergy called the Sound Life Project. Our analysis included qPCR data from one visit and CyTOF data from three longitudinal visits for each participant. Samples were collected from COVID-19 patients as soon as possible upon hospital admission, daily for the first week of hospitalization, then at 3–4-day intervals. For participants with no history of chronic disease, autoimmunity or severe allergy, samples were collected at Day 0, Day 7, and Day 90.

Both study protocols were approved by the Benaroya Research Institute Institutional Review Board (IRB20-036 for COVID-19 subjects and IRB19-045 for Sound Life Project subjects). All study procedures were conducted in accordance with the ethical standards set out in the Helsinki Declaration of 1975, as revised in 2008.

### HLA assessment via qPCR

Primers (Invitrogen/ThermoFisher) and MGB-modified probes (Applied Biosystems/ThermoFisher) for the detection of HLA-B*07 were designed for this study (Supplementary Table 1). HLA-DRA was used as an internal control (12). Reactions were amplified on an ABI StepOne Plus Sequence Detection System (Applied Biosystems, Inc). For all participants, only enrollment samples were assayed via qPCR.

### HLA assessment via CyTOF

Antibodies against HLA-A*02 and HLA-B*07 were purchased from BioLegend and BD Biosciences, respectively, and conjugated to their metal isotopes using a MaxPar X8 Multimetal Labeling Kit (Fluidigm). Antibody cocktails were made in bulk, then aliquoted and frozen prior to use as previously described (11, 13, 14). Two antibody cocktail batches were used for the samples included in this study. We did not observe a difference in staining intensity for either HLA-A*02 or HLA-B*07 between the two antibody cocktails.

Whole blood specimens were collected and stained using methods that have been previously described (11). The 33-marker panel used to identify major immune cell types has also been previously published (11) and was based on Staser et al. (15). In brief, blood was collected via venipuncture into a sterile vacutainer containing EDTA for COVID participants and sodium heparin for Sound Life Project participants. Within 24 hours of blood draw, samples were washed with phosphate-buffered saline (PBS) twice before staining with cisplatin (100 µM, Enzo Life Sciences). After a one minute room temperature incubation, cisplatin was quenched with MaxPar Cell Staining Buffer (Fluidigm). Cells were stained with thawed antibody cocktail for 20 minutes at 4°C. Red blood cell lysis was performed using RBC Lysis/Fixation solution (BioLegend) at room temperature for five minutes. Cells were washed with Cell Staining Buffer then resuspended with CELL-ID Intercalator in MaxPar Fix and Perm Buffer (Fluidigm) before storage at 4°C until data acquisition. On the day of acquisition, cells were washed with Cell Staining Buffer and Milli-Q water, then resuspended in ultrapure water containing MaxPar Four Element Calibration Beads (Fluidigm). Samples were acquired on a Helios CyTOF mass cytometer (Fluidigm) at a rate of approximately 500 events/second. We targeted 100,000 live events per sample. In all cases acquisition occurred within seven days of sample draw. The resulting data were gated manually using FlowJo software.

## RESULTS

### Calculating the percentage of errors that are detectable using two HLA antibodies

One requirement of this technique was the ability to sufficiently distinguish individuals from one another in order to enable detection of sample mismatches. We targeted a minimum of 50% detection of sample mismatches, similar to the detection ability of sex-based methods frequently used in cross-sectional studies of genomics data. To understand how well HLA-A*02 and HLA-B*07 would distinguish individuals, we estimated the percentage of study participants who would be expected to bear each HLA allele combination in our study population. That percentage allowed us to calculate the probability that, in the event of a sample swap, the two participants involved would have different HLA types, making the swap detectable via this method.

We first estimated the frequencies of HLA-A*02 and HLA-B*07 in White, Black, and Asian participants, who together make up over 90% of the population in the geographic area from which participants were recruited (16), using previously published data collected from bone marrow donors (Table 1) (17). We used an average, weighted by the racial demographics of King County, WA, USA (14), to calculate the predicted frequency of each HLA allele. We assumed our study population’s demographics would mirror those of the broader population and estimated the probability of detecting a sample swap in our population would be 51.5%. In other words, there was a 51.5% probability that samples collected from any two participants in our population would have a different combination of HLA-A*02 and HLA-B*07. This was sufficient to suggest that HLA-A*02 and HLA-B*07 were likely, in combination, to meet our goal of 50% detection of sample mismatches.

**Table 1.** Frequency (%) of HLA-A*02 and HLA-B*07 in North American Population.

| **Race** | **HLA-A\*02-**<br>**HLA-B\*07-** | **HLA-A\*02+**<br>**HLA-B\*07-** | **HLA-A\*02-**<br>**HLA-B\*07+** | **HLA-A\*02+**<br>**HLA-B\*07+** | **Probability of**<br>**detecting a swap (%)** |
| --- | --- | --- | --- | --- | --- |
| Asian* | 71.1 | 24.2 | 3.8 | 0.5 | 43.0 |
| Black, African American* | 70.5 | 17.4 | 9.1 | 1.5 | 44.9 |
| White, Caucasian* | 59.2 | 25.2 | 8.7 | 3.4 | 54.3 |
| Total Cohort | 61.4 | 24.8 | 8.0 | 2.9 | 51.5 |

\*Based on published allele frequencies(17)
\*Excluding participants of unknown or mixed race from the probability assessment
Note: frequencies do not sum to 100% due to rounding

We included 41 participants in our study. The racial make-up of the cohort can be seen in Supplementary Table 2 and was generally representative of the population of King County, WA. Using the actual racial make-up of our cohort, we calculated that our method had a 52.5% probability of correctly identifying sample swaps and a 47.5% probability of missing a swap because it occurred between participants with the same combination of HLA-A*02 and HLA-B*07 alleles.

### HLA-A*02 and HLA-B*07 expression varies by cell population

HLA-A*02 and HLA-B*07 specific antibodies were incorporated into a CyTOF panel designed to identify major immune cell populations. Our gating strategy was previously used to characterize major immune cell populations in whole blood collected from SARS-CoV-2+ patients (11). To help inform our strategy for identifying participants who were positive for either HLA allele, we examined median metal intensity in 10 immune cell populations, including the overall CD45+ population (Figure 1). We observed low expression levels of both alleles in the neutrophil population, which likely contributed to low expression levels observed in the overall CD45+ population. We observed higher expression of both alleles among mononuclear cell subsets, particularly monocytes, B cells, and NK cells. Overall, staining for HLA-B*07 was less intense than HLA-A*02 in our assay. To ensure that we were gating on a sufficiently large population of cells and to reduce false negatives caused by low HLA expression, we chose to gate for HLA-A*02 and HLA-B*07 positive cells among non-granulocytes—the general CD45+ population with the granulocytes (basophils, eosinophils, and neutrophils) gated out. This selection also allows this technique to be readily transferrable to frozen PBMC samples, which lack these polymorphonuclear subsets.

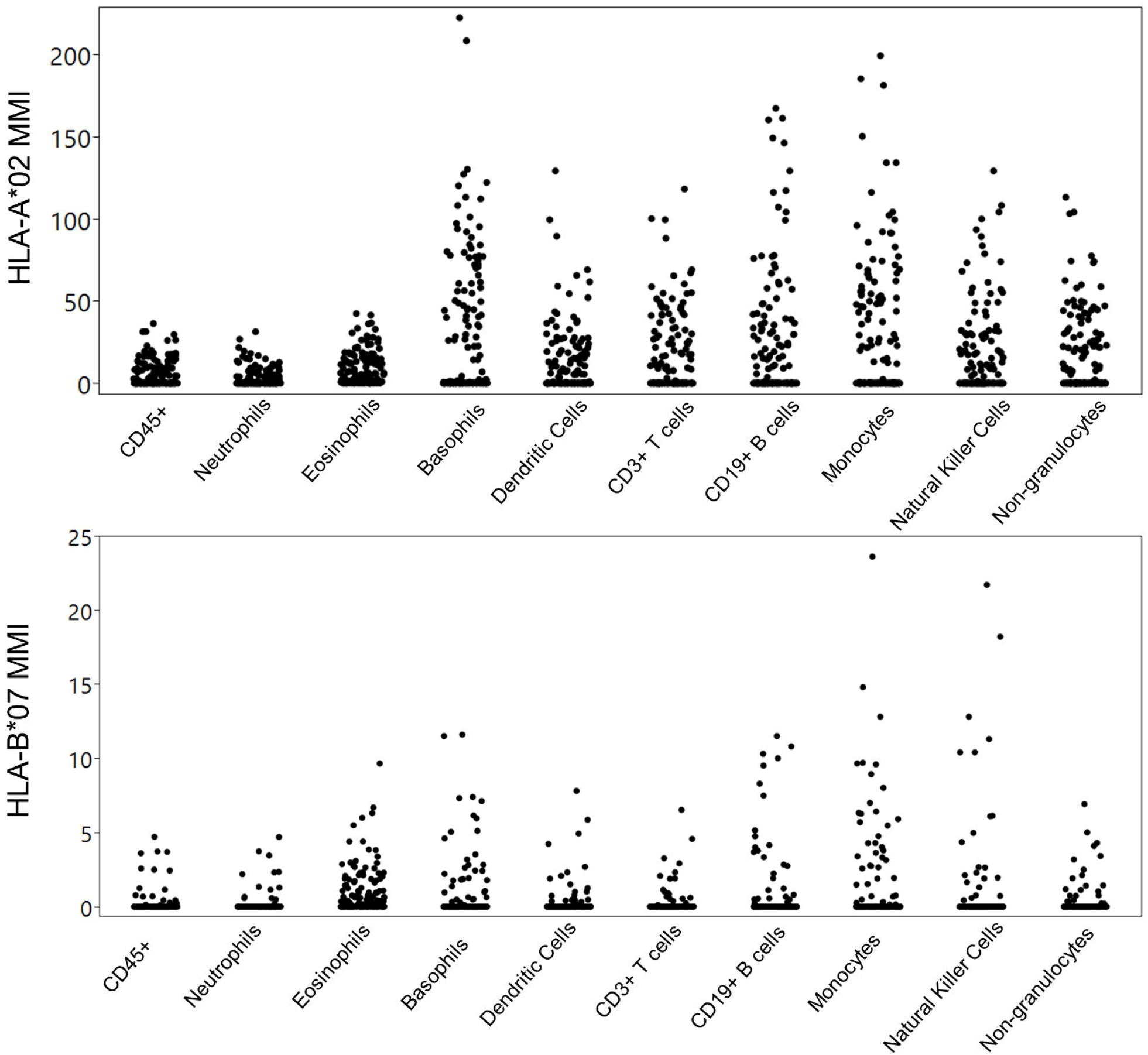

Representative gating in participants who were positive and negative for each allele is shown in Figure 2. Samples were considered positive for the HLA-A*02 allele if the frequency of parent was above 20%. Samples were considered positive for the HLA-B*07 allele if the frequency of parent was above 5%.

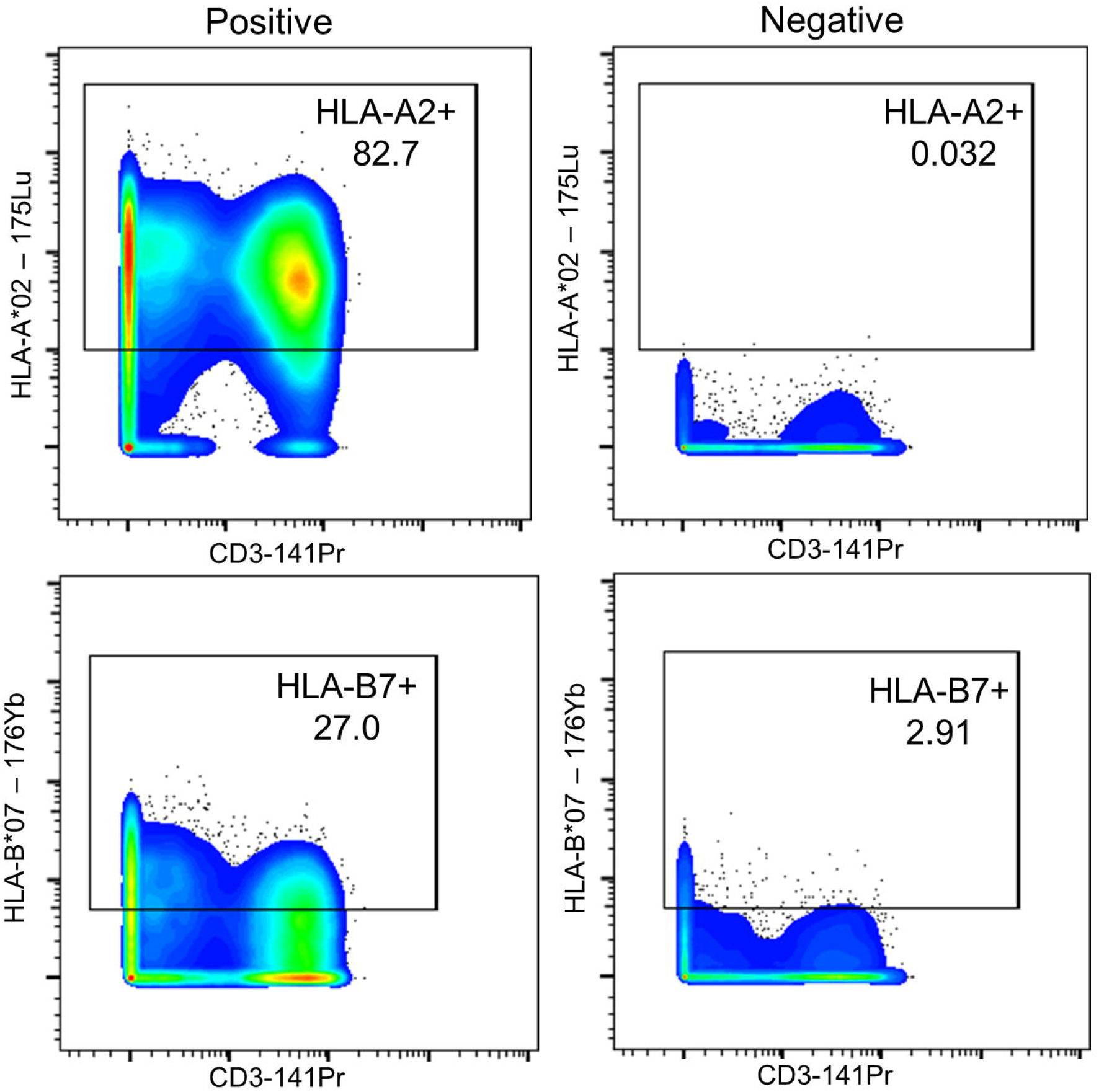

### Identifying sample annotation errors

We classified each sample as positive or negative for HLA-A*02 and HLA-B*07, and compared these classifications taken from CyTOF data at three timepoints. We identified three longitudinal sample sets with a case of mismatching allele expression (Figure 3). In all three cases, the identified mismatch was in the HLA-B*07 cytometry data; no mismatches for HLA-A*02 were identified. In two cases, the apparent mismatched sample was negative for HLA-B*07 while the other 2 samples were positive for this allele. For one subject, the apparent mismatched sample was positive while other samples were negative. qPCR data for HLA-B*07 agreed with the majority of the cytometry data, with Participants 1 and 2 being positive and Participant 3 being negative for HLA-B*07.

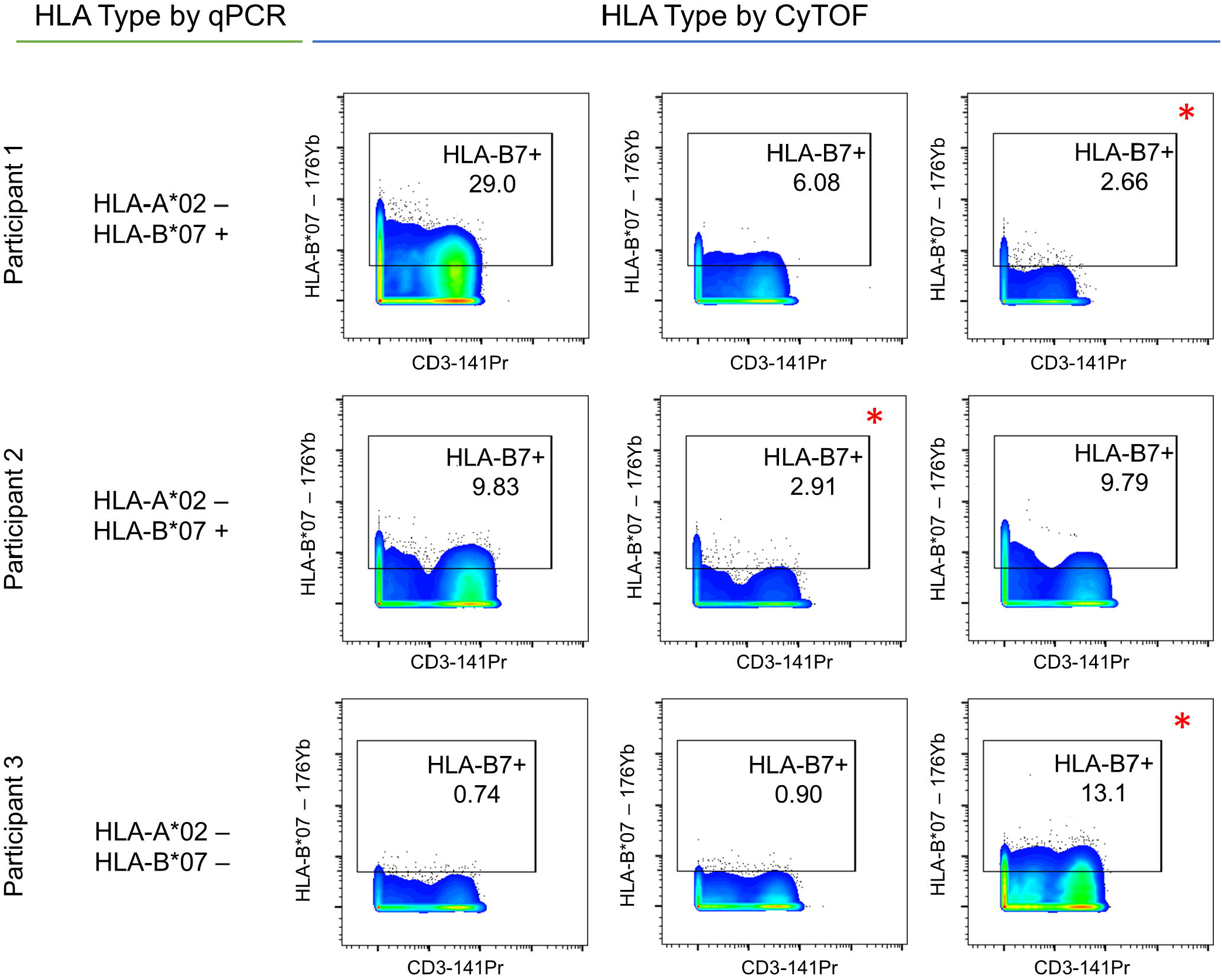

### Sample annotation error rate estimation

Comparison of longitudinal CyTOF data allowed us to identify three samples that were paired with the incorrect metadata; this was confirmed by qPCR. These three samples constitute 2.4% (3/123) of all the CyTOF samples included in this analysis. As this method is predicted to detect approximately half of all mismatches in our cohort, the true error rate is likely approximately 5%. This error rate is similar to the error rate reported by others investigating publicly available genomic datasets (1, 6) using methods with similar detection levels.

## DISCUSSION

We developed a technique that allows investigators to perform quality control on cytometry datasets by measuring HLA class I alleles. We demonstrate that this technique can be used in longitudinal samples to identify sample annotation errors in cytometry data. We also show this technique can be used in conjunction with a secondary method, like qPCR, which could be implemented to assess data validity in cross-sectional studies when longitudinal data is unavailable. This novel technique could aid investigators performing cytometry on clinical samples in avoiding the commonly identified problem of misannotation (1, 2).

When choosing HLA class I alleles as our target proteins, we considered several important characteristics. For a clinical study of human blood samples, our target proteins needed to be: detectable by cytometry, consistently expressed on blood cells, and measurable with commercially available reagents. The medical community has a longstanding interest in HLA typing because of its importance in assessing compatibility of organ donors (18), creating an incentive to develop a variety of typing techniques and commercially available reagents. Indeed, a technique for HLA typing of subjects via flow cytometry has already been published (19), and HLA typing for selection of samples in tetramer assays has been in place for some time. Our target proteins needed to be detectable over time with expression remaining robust during immune challenges like disease or vaccination. HLA class I proteins are expressed on the surface of most nucleated cells (18). Even in cases where cell subset frequencies fluctuate, such as COVID-19 infection, a large number of HLA class I presenting cells remain present in the sample.

Additionally, our target proteins needed to be sufficiently unique to individual participants. The MHC is the most highly polymorphic region of the human genome with over 25,000 alleles (20), providing us with many targets to choose from. We chose two HLA alleles HLA-A*02 and HLA-B*07 which together gave us a > 50% probability that any two samples in our dataset would be distinguishable, a level of detectability that is similar to existing methods for sex-based detection of misannotation.

Investigators wishing to maximize their ability to detect swaps could consider increasing the number of target proteins included in their panel. Other potential target proteins include additional HLA class I alleles, HLA class II alleles, or other polymorphic proteins like FcgRIII or KIR. Alternatively, targets that are differentially expressed by sex, such as H-Y antigen or HDH1, KDM6A, EIF1AX, UTY, DDX3Y, and ZFY could be appropriate. When choosing a target, investigators should be aware of which cell subsets express target proteins and how those cell subsets vary over the course of the study. Investigators may also consider choosing target proteins that are as evenly distributed across their population as possible. For example, a panel including two target proteins, Allele A and Allele B, will have maximum detectability (75%) if 25% of the population is double positive, 25% is positive for Allele A and negative for Allele B, 25% is negative for Allele A and positive for Allele B, and 25% is double negative. As the allele distribution becomes less even, the detectability will also decrease. In this study, the addition of a sex chromosome-associated marker, which would have been evenly distributed across our cohort, would have increased the probability that we would detect a swap to 74.6%. In all studies, the ability to add target proteins to detect a swap must be balanced against the need to answer the primary study question.

In this study, we used qPCR as a secondary measure to validate HLA-type when HLA was already classified using CyTOF, but similar genomic methods could also be used as an external data source for quality control of cross-sectional data when longitudinal data is unavailable. Investigators should be aware, however, that if genomic and cytometric data are discordant, it may be difficult to determine where the error occurred, and both sets of data may need to be excluded from analysis or further investigated.

For the analysis of longitudinal cytometry data, we found this technique was useful for identifying potentially misannotated samples. However, we did identify several key limitations. This technique was not designed to identify sample swaps between individuals of the same HLA-type. This limitation is likely to be shared, to varying degrees, by other polymorphic or sex-associated protein targets. Another limitation of this technique was the potential for non-specific binding of the HLA antibodies. This limitation could cause samples that are correctly annotated to be identified as erroneous. We also note that the HLA-B*07 frequency of parent determined by CyTOF was close to our chosen cut off for HLA-B*07 positivity in some apparent mismatched samples. Given the low expression of HLA-B*07 in all samples, and the variability we observed in staining intensity, it is likely that our HLA-B*07 antibody was not saturating. We would recommend careful titration of the antibody to others wishing to utilize this technique. In this case, confirmation of the HLA type by qPCR provided further evidence that these three samples were misannotated.

This technique – with antibody modifications specific to any given study - is a novel and important tool for ensuring the integrity of cytometry datasets. As a community we must accelerate the development of sample identification approaches so that mismatches between assay and clinical data can be detected when they occur. Incorporating this quality control will ensure that scientific advances based on cytometry data are accurate and reproducible.

## Supporting information

Supplemental Table 1. PCR Primers and Probes

Supplemental Table 2. Participant Demographics and Clinical Characteristics

## Acknowledgements

We would like to thank Uma Malhotra and the Benaroya Research Institute (BRI) COVID-19 Research Team for their help in obtaining COVID-19 participant samples and data; Kassidy Benoscek, Deric Khuat, and Claire Mangan for Sound Life Project study recruitment and conduct; Thien-Son Nguyen and the BRI Clinical Core Laboratory for sample processing; and Rachel Hartley for data management. Thanks to Colin O’Rourke for consulting on allele detection probability calculations. Special thanks to Nicole Gilbert for generating the qPCR data. Thanks to Kerry Casey for assistance in the design of the CyTOF panel. JHB is a Scientific Co-Founder and Scientific Advisory Board member of GentiBio, a consultant for Bristol-Myers Squibb and Hotspot Therapeutics, and has past and current research projects sponsored by Amgen, Bristol-Myers Squib, Janssen, Novo Nordisk, and Pfizer. She is a member of the Type 1 Diabetes TrialNet Study Group, a partner of the Allen Institute for Immunology and a member of the Scientific Advisory Boards for the La Jolla Institute for Allergy and Immunology and BMS Immunology. JHB also has a patent on methods for generating antigen-specific CD4+CD25+ regulatory T cells.

---

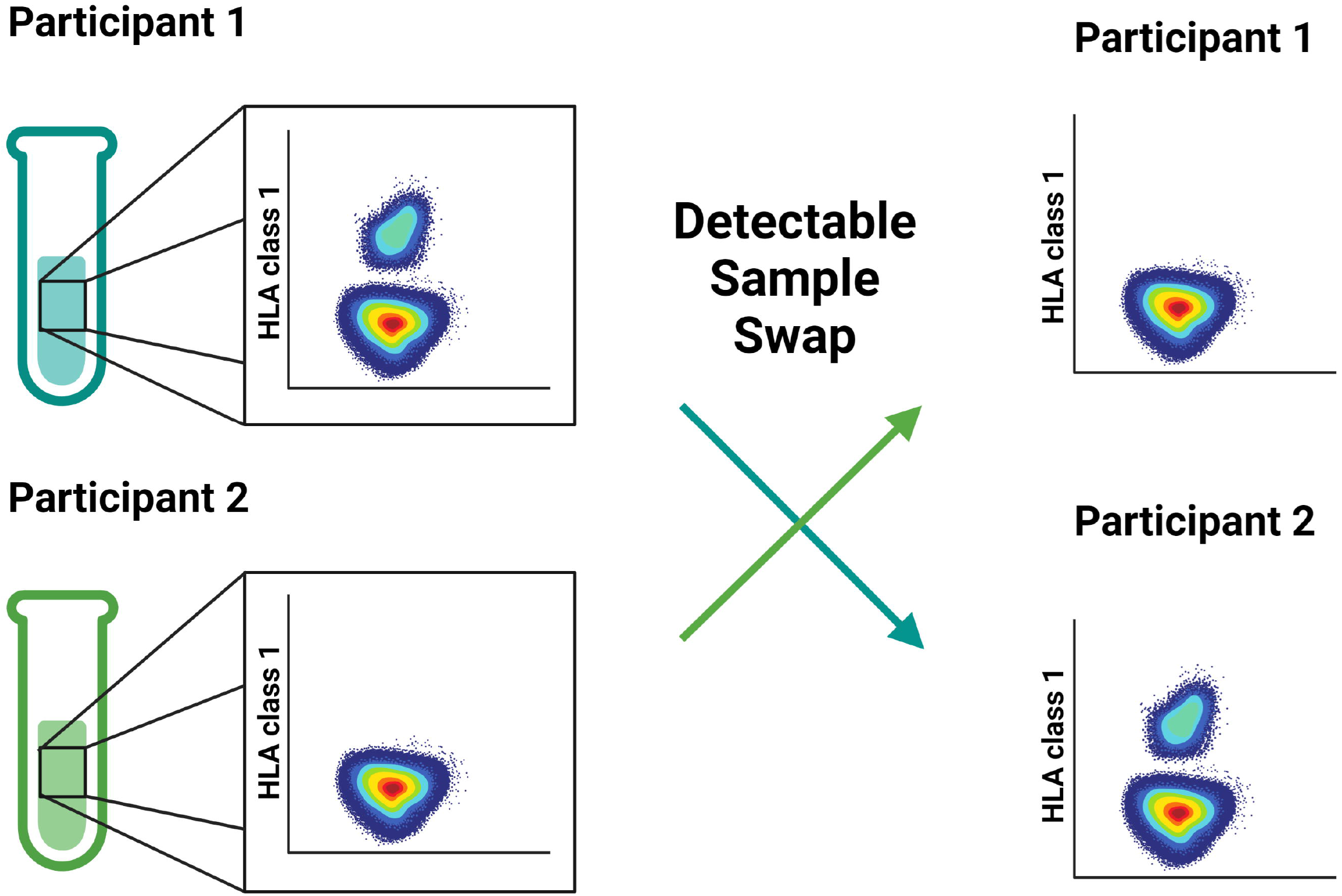

## Notes

**Funding:** The Allen Institute for Immunology provided funding for the development of the CyTOF panel and for recruitment and sample analysis of the healthy control subjects. The Benaroya Family Foundation, the Leonard and Norma Klorfine Foundation, and Glenn and Mary Lynn Mounger supported the COVID-19 studies.

**Conflict of Interest:** JHB is a Scientific Co-Founder and Scientific Advisory Board member of GentiBio, a consultant for Bristol-Myers Squibb and Hotspot Therapeutics, and has past and current research projects sponsored by Amgen, Bristol-Myers Squib, Janssen, Novo Nordisk, and Pfizer. She is a member of the Type 1 Diabetes Trialnet Study Group, a partner of the Allen Institute for Immunology and a member of the Scientific Advisory Boards for the La Jolla Institute for Allergy and Immunology and BMS Immunology. JHB also has a patent on methods for generating antigen-specific CD4+CD25+ regulatory T cells.

