## Supplemental Table 1. PCR Primers and Probes for "A simple strategy for sample annotation error detection in cytometry datasets"

**Supplementary Table 1. PCR primers and probes**

| **Allele** | **Forward Primer** | **Reverse Primer** | **Probe** | **Reference** |
| --- | --- | --- | --- | --- |
| HLA-B*07 | AGATCACCCAGCGCAAGTG | GGAGCCACTCCACGCACTC | AGCAGCGGAGAGCC | Developed in-house |
| HLA-DRA | AGGCCGAGTTCTATCTGAATCCT | CGCCAGACCGTCTCCTTCT | CATAAACTCGCCTGATTG | (1) |
| 1. Gersuk VH, Nepom GT. A real-time PCR approach for rapid high resolution subtyping of HLA-DRB1*04. Journal of Immunological Methods. 2006;317:64-70. | | | |  |
