## Supplemental Table 2. Participant Demographics and Clinical Characteristics for "A simple strategy for sample annotation error detection in cytometry datasets"

**Supplementary Table 2. Participant demographics and clinical characteristics**

| **Population Characteristics** | **n (%)** |
| --- | --- |
| Disease State |  |
| Healthy Control | 25 (61.0) |
| COVID-19 | 16 (39.0) |
| Race |  |
| Asian | 5 (12.2) |
| Black, African American | 2 (4.9) |
| White, Caucasian | 30 (73.2) |
| Mixed Race or Other Race | 4 (9.8) |
| Female | 23 (56.1) |
| Age, median (range) | 58 (27 - 89) |
